# Carriage of antibiotic resistant bacteria in endangered and declining Australian pinniped pups

**DOI:** 10.1101/2021.10.11.463979

**Authors:** Mariel Fulham, Fiona McDougall, Michelle Power, Rebecca R. McIntosh, Rachael Gray

## Abstract

The rapid emergence of antimicrobial resistance (AMR) is a major concern for wildlife and ecosystem health globally. Genetic determinants of AMR have become indicators of anthropogenic pollution due to their greater association with humans and rarer presence in environments less affected by humans. The objective of this study was to determine the distribution and frequency of the class 1 integron, a genetic determinant of AMR, in both the faecal microbiome and in *Escherichia coli* isolated from neonates of three pinniped species. Australian sea lion (*Neophoca cinerea*), Australian fur seal (*Arctocephalus pusillus doriferus*) and long-nosed fur seal (*Arctocephalus forsteri*) pups from eight breeding colonies along the Southern Australian coast were sampled between 2016-2019. DNA from faecal samples (*n*=309) and from *E*. *coli* (*n*=795) isolated from 884 faecal samples were analysed for class 1 integrons using PCRs targeting the conserved integrase gene (*intI*) and the gene cassette array. Class 1 integrons were detected in *A*. *p*. *doriferus* and *N*. *cinerea* pups sampled at seven of the eight breeding colonies investigated in 4.85% of faecal samples (*n*=15) and 4.52% of *E*. *coli* isolates (*n*=36). Integrons were not detected in any *A*. *forsteri* samples. DNA sequencing of the class 1 integron gene cassette array identified diverse genes conferring resistance to four antibiotic classes. The relationship between class 1 integron carriage and the concentration of five trace elements and heavy metals was also investigated, finding no significant association. The results of this study add to the growing evidence of the extent to which antimicrobial resistant bacteria are polluting the marine environment. As AMR determinants are frequently associated with bacterial pathogens, their occurrence suggests that these pinniped species are vulnerable to potential health risks. The implications for individual and population health as a consequence of AMR carriage is a critical component of ongoing health investigations.

## Introduction

Aquatic ecosystems are being increasingly identified as a sink for antimicrobial resistance (AMR) (1,2). Aquatic systems provide a transport medium for the global dissemination of antibiotic resistant bacteria (ARB) and associated antibiotic resistance genes (ARGs) (1–3). The combination of ARB with antibiotic residues and other pollutants in aquatic environments also promote the proliferation and establishment of resistant bacterial communities (2,4).

The widespread dissemination of AMR can partly be attributed to horizontal gene transfer (HGT) which allows the transfer of ARGs and associated genetic machinery between diverse bacterial species, facilitating the acquisition of novel traits from the environment and other bacteria (5,6). In Gram-negative bacteria in particular, the rapid evolution of resistance has been linked to HGT and mobile genetic elements (7). Class 1 integrons, for example, are mainly found in Gram-negative bacteria (8) and are able to capture and subsequently express a multitude of ARGs (9,10), which can be transferred between bacteria via their association with transposons and plasmids. In the human context, the clinical class 1 integron is considered to be of high importance for AMR dissemination (11). The class 1 integron contains a conserved 5’ segment which encodes the integrase gene (*intI1*) and a varying number of gene cassettes that together form a gene cassette array (12). The conserved *intI1* is a useful genetic indicator of antimicrobial pollution as it is universally present, occurs in high abundance in humans and domestic animals, is highly abundant in waste streams and is rarely present in environments less affected by humans (13). The recombination of gene cassettes is mediated by *intI1*, allowing the class 1 integron to capture, remove and express a variety of gene cassettes (12). Variations of the class 1 integron are now emerging, with insertion sequences in the 3’ segment, such as IS*26*, assisting in the dissemination of resistance genes in Gram-negative bacteria. These insertion sequences are associated with numerous genes that confer resistance to multiple antibiotic classes, and are able to promote and subsequently express these associated resistance genes (14,15).

Agricultural runoff, in addition to mining, municipal wastewater, and industrial and pharmaceutical waste are point sources of heavy metal pollutants frequently found in natural environments (16). Heavy metals are considered to be co-selective agents of AMR (16). Aquatic environments polluted by heavy metals have been associated with a greater incidence of class 1 integrons compared to non-polluted sites (17,18), through mechanisms of cross- and co-resistance (19). The presence of heavy metals and antibiotic residues also common in the environment have the potential to exert a selective pressure that promotes the emergence and persistence of AMR in the environment (20–22). As heavy metals can bioaccumulate and persist in the environment, such selective pressures are applied for extended periods of time, facilitating the development resistance traits in microbial communities (23). However, there has been little investigation into whether there is increased acquisition of antibiotic resistant bacteria in humans and non-human animals in environments that have greater exposure to heavy metals.

Concentrations of essential trace elements and heavy metals including zinc (Zn), arsenic (As), selenium (Se), mercury (Hg) and lead (Pb) are of particular interest for wildlife health. The presence of Pb, even at low concentrations, can be associated with disease (24). In contrast, Zn and Se are essential trace elements, but these too can have toxic effects at high concentrations (25). Heavy metals have previously been identified in free-ranging pinnipeds (26–28), however, there has been no consideration of potential co-selection of ARGs in wildlife species associated with heavy metal exposure. Given the role that heavy metals play in the environmental amplification of ARB, investigating the levels of heavy metals and class 1 integron frequency in wildlife species could provide valuable insights into the factors contributing to the abundance and acquisition of ARB in free-ranging wildlife.

A diverse range of antibiotic resistant bacterial species have been detected in marine mammals (29,30), which are large-bodied, long-lived upper trophic predators considered to have a role as sentinels of marine health (31). *Escherichia coli* is a Gram-negative bacterium that is commonly used as an indicator of anthropogenic pollution (20), and has been used for the investigation of class 1 integrons in many species (32–36). Evidence suggests that class 1 integrons are more prevalent in *E*. *coli* isolates that are closely associated with anthropogenic pollution and human environments (7). The presence of class 1 integrons in *E*. *coli* has been investigated in some species of free-ranging pinnipeds in the Southern Hemisphere. An absence of class 1 integrons was reported in *E*. *coli* isolated from free-ranging southern elephant seals (*Mirounga leonina*), Weddell seals (*Leptonychotes weddellii*) (37) and adult Australian sea lions (*Neophoca cinerea*) (36), although class 1 integrons were detected in *E*. *coli* from captive adult *N*. *cinerea* (36). The presence of class 1 integrons in captive wildlife and comparative absence in free-ranging individuals suggest that environmental conditions and the intimate proximity to humans experienced in captivity can impact the acquisition of ARGs by wildlife species and is consistent for many wildlife species (33,34,36). Consistent with the hypothesis that the presence of ARGs in wildlife is associated with proximity to humans, a class 1 integron was recently detected in *E*. *coli* from a single free-ranging *N*. *cinerea* pup at a colony with comparatively high anthropogenic influence compared to a more remote colony (35).

The occurrence of class 1 integrons and ARG carriage in two additional pinniped species inhabiting Australian waters, namely Australian fur seals (*Arctocephalus pusillus doriferus*) and long-nosed fur seals (*Arctocephalus forsteri*), has not been investigated. All three species, *N*. *cinerea*, *A*. *p*. *doriferus*, and *A*. *forsteri*, inhabit numerous offshore islands along the Australian coast from Western Australia to Tasmania (38), with the ranges of these pinniped species overlapping in South Australia. Colonies of each species experience differing levels of human interaction; those on islands remote to mainland Australia experience little to no contact with humans while others are popular, frequently visited tourist sites. The differing proximities of sympatric colonies to human habitation and exposure to anthropogenic impacts creates a naturally occurring gradient ideal for studying anthropogenic pollution in the marine environment.

The main objective of this study was to determine the prevalence of class 1 integrons and ARG carriage in both the faecal microbiota and *E*. *coli* isolates from pups of three pinniped species sampled at multiple breeding colonies throughout Southern Australia. Given the role heavy metals have as a co-selective agent for AMR, an additional aim was to determine whether there was a relationship between the concentration of essential elements and heavy metals (Zn, Se, As, Hg and Pb) and class 1 integron prevalence. It was hypothesised that class 1 integrons would be more abundant in pups at colonies in closer proximity to sources of anthropogenic pollution. This paper reports the presence of ARGs in *E*. *coli* isolates and faecal microbiota of *N*. *cinerea* and *A*. *p*. *doriferus* pups at multiple breeding colonies along the Australian coast. We explore the differences between species and colonies and discuss factors contributing to changes in class 1 integron prevalence across breeding seasons and colonies. We provide recommendations for future investigations to further understand the dissemination of AMR in free-ranging pinniped species.

## Methods

### Study sites and sample collection

Faecal swabs (*n*=884) were collected from neonatal pups sampled at eight breeding colonies across multiple breeding seasons from 2016-2019 (Fig 1 and Table 1). Breeding seasons are annual for both *A*. *p*. *doriferus* and *A*. *forsteri* with pupping beginning in November, while *N*. *cinerea* breeding seasons occur every 18 months. *Arctocephalus pusillus doriferus* and *A*. *forsteri* pups were approximately 3-6 weeks of age and *N*. *cinerea* pups were 2-6 weeks of age at time of sampling. Samples were collected following methods described by Fulham et al. (35,39). In brief, sterile swabs (Copan, Brescia, Italy) were inserted directly into the rectum of each pup and resulting samples were sub-sampled into sterile FecalSwab™ tubes (Copan, Brescia, Italy). FecalSwab samples were stored at 4°C and cultured within 7-10 days of collection. Blood samples were collected from the brachial vein of pups as per methodology in Fulham et al. (35) and refrigerated at 4°C prior to storage at −20°C. Due to time and logistical constraints, blood collection was limited to pups sampled at Seal Bay, Olive Island, Cape Gantheaume, Seal Rocks (2018 and 2019) and Deen Maar Island. Sampling for *N*. *cinerea* and *A*. *forsteri* were approved by the Animal Ethics Committee at the University of Sydney (Protocol Nos. 2014/726 and 2017/1260); sampling methods for *A*. *p*. *doriferus* were approved by Phillip Island Nature Parks Animal Ethics Committee (Protocol No. 2.2016).

**Fig 1.**
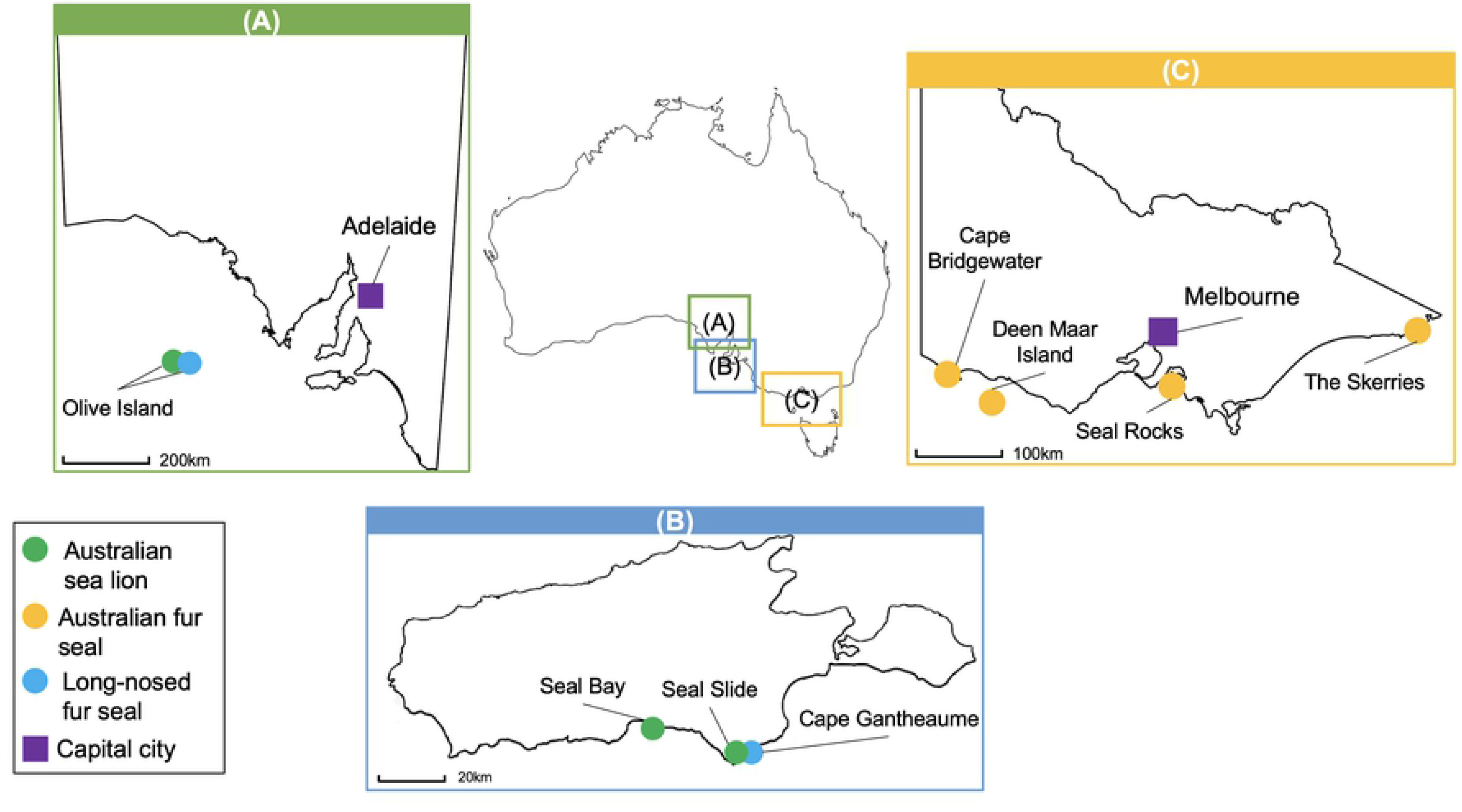
Map of the geographical locations of breeding colonies and pinniped species sampled. Breeding colonies in South Australia (A) include Olive Island, (B) Seal Bay, Seal Slide and Cape Gantheaume on Kangaroo Island, and (C) Cape Bridgewater, Deen Maar Island, Seal Rocks and The Skerries in Victoria. The closest capital city to breeding colonies in South Australia is Adelaide and capital city in Victoria is Melbourne.

**Table 1.** Sample collection across breeding colonies and seasons.

| Sampling site | Species | Year of breeding<br>season and sample<br>collection (*n* faecal<br>samples collected) | Geographical<br>coordinates |
| --- | --- | --- | --- |
| Seal Bay | *N. cinerea* | 2016 (48) | 35.95°S, 137.32°E |
|  |  | 2018 (59) |  |
|  |  | 2019 (63) |  |
| Seal Slide | *N. cinerea* | 2018 (4) | 36.03°S, 137.29°E |
| Olive Island | *N. cinerea* | 2017 (89) | 32.43°S, 133.58°E |
|  |  | 2019 (66) |  |
|  | *A. forsteri* | 2019 (12) |  |
| Cape Gantheaume | *A. forsteri* | 2016 (69) | 36.24°S, 137.27°E |
|  |  | 2018 (80) |  |
| Seal Rocks | *A. p. doriferus* | 2017 (46) | 38.31°S, 145.5°E |
|  |  | 2018 (99) |  |
|  |  | 2019 (94) |  |
| Deen Maar Island | *A. p. doriferus* | 2017 (95) | 38.24°S, 142.0°E |
| Cape Bridgewater | *A. p. doriferus* | 2017 (37) | 38.18°S, 141.24°E |
| The Skerries | *A. p. doriferus* | 2017 (23) | 37.45°S, 149.31°E |

Location of sampling sites, species at each colony and number of faecal samples collected during each breeding season.

### *E*. *coli* culture, isolation and DNA extraction

FecalSwab™ samples were cultured following the methodology described by Fulham et al. (35,39). In summary, a selective media, Chromocult® coliform agar (Merck, Darmstadt, Germany), was used to isolate *E*. *coli*. After initial culture, *E*. *coli* colonies were sub-cultured and pure *E*. *coli* colonies were selected for DNA extraction based on colour and morphology. Pure *E*. *coli* colonies were inoculated into Luria-Bertani broth (5ml) and incubated at 37°C for 24 hours in preparation for preservation and DNA extraction. DNA was extracted using a boil preparation method where the broth culture was centrifuged to pellet bacteria. Supernatant was decanted and bacterial pellet resuspended with sterile water (50μL). Samples were then heated for 5min at 95°C followed by centrifugation and resulting bacterial lysates were stored at −30°C.

### Faecal DNA extraction and PCR competency

DNA was extracted from a subset of faecal samples (*n*=309) as part of an exploratory analysis into the presence of *intI1* in pinniped microbiomes. Samples were randomly selected from each of the following colonies during each breeding season: Seal Bay 2016 (*n*=48); Seal Rocks 2017 (*n*=46), 2018 (*n*=30), 2019 (*n*=30); Deen Maar Island (*n*=30); Cape Bridgewater (*n*=30); the Skerries (*n*=23); Cape Gantheaume 2018 (*n*=30); Olive Island 2019 (*n*=30 *N*. *cinerea*, *n*=12 *A*. *forsteri*). Genomic DNA was extracted from FecalSwab™ sample media (200μL) using the ISOLATE Fecal DNA kit (Bioline, Sydney, Australia) as per manufacturer’s instructions. PCR competency of DNA extracted from faecal samples and *E*. *coli* isolates was tested by a 16S PCR (Table 2) using methods described by Fulham et al. (35).

**Table 2.**
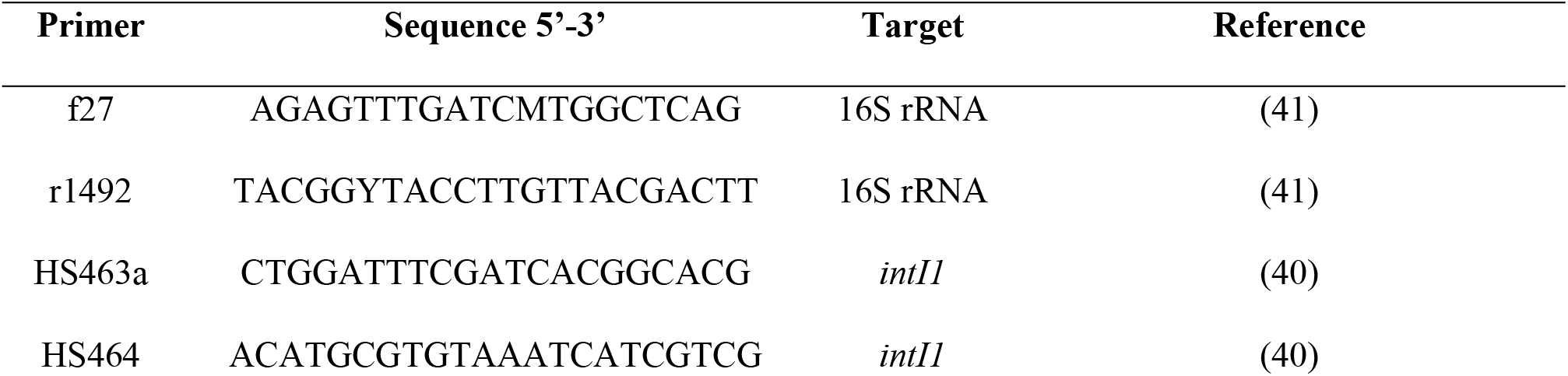
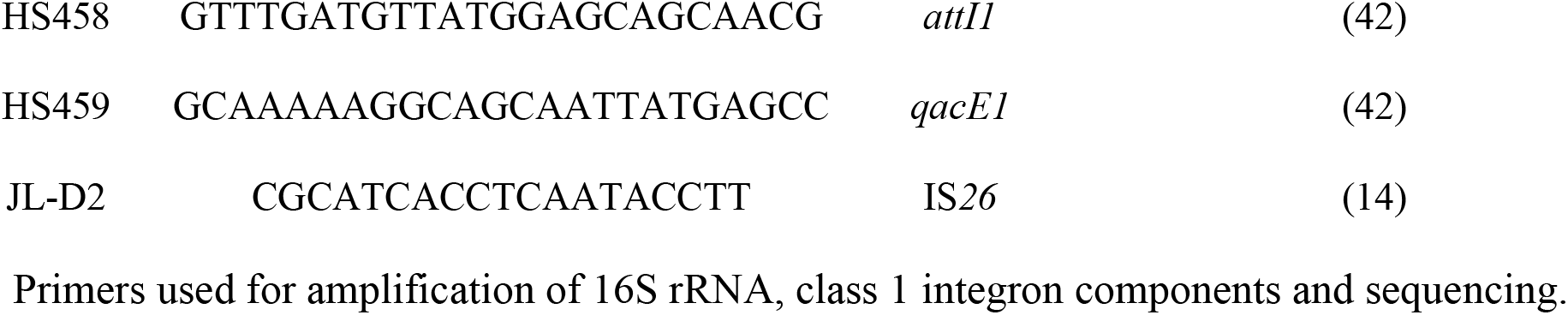
PCR primers and target region.

### Screening for class 1 integrons

All faecal samples and *E*. *coli* isolates positive for 16S rDNA were further screened for the presence of the class 1 integron integrase gene (*intI1*) using HS463a and HS464 primers (Table 2) following the methods described by Waldron and Gillings (40).

Samples containing *intI1* were tested using additional PCRs to target the gene cassette array using HS458 and HS459 primers (Table 2) and PCR conditions described by (40). Any samples that did not produce a band for the HS458/HS459 PCR were analysed using a secondary primer set consisting of HS458 and JL-D2, which targets the IS*26* transposase, an alternate 3’ terminus in integrons (14), using the conditions as described for HS458/HS459.

All PCRs included a positive control sample (integron positive *E*. *coli* KC2) and negative control (PCR-grade H_2_O) and were resolved using gel electrophoresis (16S rRNA and HS463a/HS464 2% agarose w/v, HS458/459 and HS458/JL-D2 3% agarose w/v) with SYBR safe gel stain (Invitrogen, city, Australia). Electrophoresis was conducted at 100V for 30 min (16S) or 40 min (463/464; 458/459/JL-D2) in TBE (Tris, boric acid, ethylenediaminetetraacetic acid) and product size approximated using HyperLadderII 50bp DNA marker (Bioline, Sydney, Australia).

### Cloning, sequencing and analysis

Using the MinElute PCR Purification Kit (Qiagen, Melbourne, Australia), amplicons from the two gene cassette array PCRs (HS458/459 and HS458/JL-D2) were purified following manufacturer’s instructions. Amplicons containing only a single band were sequenced directly using the purified PCR product.

Amplicons containing multiple bands, indicating the presence of more than one gene cassette, were cloned using the TOPO TA cloning kit and transformed into One Shot ® DH5™-T1^R^ competent *E*. *coli* cells (Invitrogen, Carlsbad, CA, USA) as per manufacturer’s protocol. Between six to twelve colonies of transformed *E*. *coli* were selected from each cloned sample and DNA from cell lysates were screened using HS458, HS459 and JL-D2 as described above. Amplicons of variable sizes were selected for sequencing. Amplicons from HS463a/HS464 that did not amplify in HS458/HS459 or HS458/ JL-D2 were purified and sequenced to confirm positive *intI* result.

Sequencing was performed at The Ramaciotti Centre for Genomics (University of New South Wales, Sydney, Australia) using Big Dye Terminator chemistry version 3.1 and ABI 3730/3730×1 Capillary Sequencers (Applied Biosystems, Foster City, CA, USA). Geneious Prime software (version 11.0.6; Biomatters Limited, Auckland, New Zealand) was employed to assemble and manually check sequences for quality. Assembled sequences were analysed for the presence of antibiotic resistance genes using Integrall (http:/integrall.bio.ua.pt). Class 1 integron gene cassette arrays were confirmed via detection of the 3’ conserved region containing *qacE*. For arrays containing more than one gene cassette, the *attC* recombination site located between cassettes was identified using the highly conserved core sequence GTTRRRY and complementary inverse core sequence RYYAAC (43). Representative sequences generated from this study have been submitted to GenBank and are awaiting accession.

### Essential element and heavy metal concentrations

The concentrations of Zn, As, Se, Hg, and Pb in whole blood of *A*. *p*. *doriferus* pups sampled at Seal Rocks in 2018 (*n*=52) were provided by another study (Cobb-Clarke and Gray, personal communication) following previously described methods (27). The data was derived from samples analysed using inductively coupled plasma-mass spectrometry (ICP-MS; Agilent Technologies 7500 ce inductively coupled plasma mass spectroscopy, Santa Clara, CA). The median values and 95% confidence intervals (obtained from back transformed log data) for essential element and heavy metals in blood (μg/L) were Zn=3.73 (95% CI 3.67-3.87), Se=3.05 (95% CI 3.00-3.56), As=0.06 (95% CI 0.05-0.07), Hg=0.04 (95% CI 0.05-0.12) and Pb=0.04 (95% CI 0.04-0.10).

### Statistical analyses

All statistical analyses were conducted using RStudio software (V 1.2.5042, Boston, Massachusetts, USA). The Shapiro-Wilk’s test was used to test for normality of data and any variables with significant (<0.05) and non-normal distribution were log transformed which normalised the data set and allowed for parametric statistical analysis. Significance was determined when p < 0.05.

The statistical analysis of class 1 integron distribution was conducted using Fisher’s exact test to test for differences in class 1 integron occurrence between species. Pearson’s chi-squared test was used to test for differences in class 1 integron occurrence within species across sampling sites and breeding seasons.

Welch’s two sample t-test was used to test for significance in the relationship between integrons and essential element and heavy metal concentrations in *A*. *p*. *doriferus* pups (*n*=52) sampled at Seal Rocks 2018. Of these 52 individuals, 14 were integron positive.

## Results

### Detection of class 1 integrons

*Escherichia coli* was isolated from a total of 795 faecal samples (89.9%) with PCR screening for *intI1* revealing 36 positive isolates (4.52%, *n*=795) based on the presence of the expected 473bp product. Seven of the positive *E*. *coli* isolates were from *N*. *cinerea* and 29 from *A*. *p*. *doriferus*. Screening of faecal DNA detected *intI1* in 15 faecal samples (4.85%, *n*=309), with four from *N*. *cinerea* and 11 from *A*. *p*. *doriferus*. All faecal samples and *E*. *coli* isolates from *A*. *forsteri* were negative for *intI1* (Fig 2).

**Fig 2.**
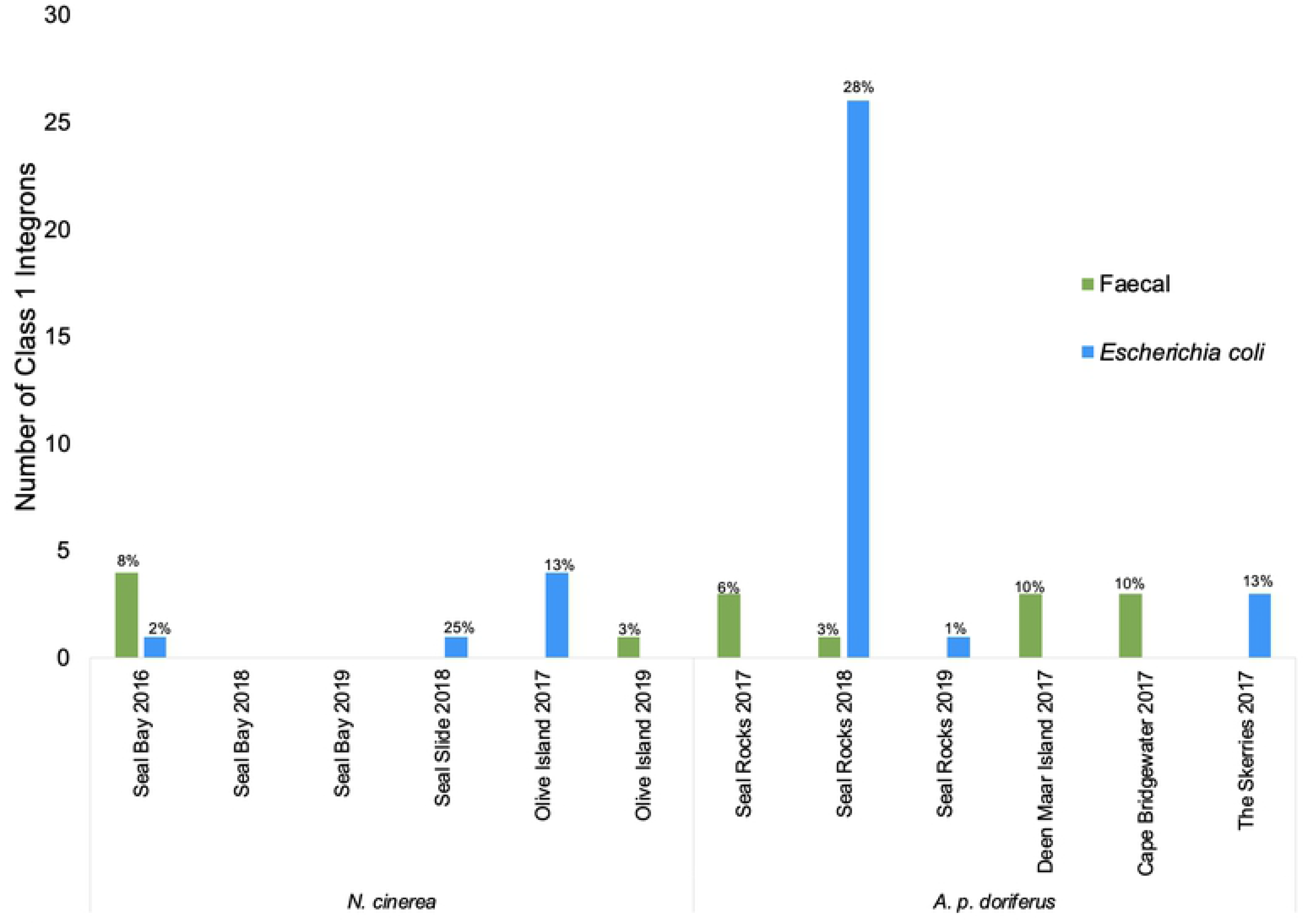
Graph of class 1 integrons detected in pinniped pups. Total number of class 1 integrons detected in faecal and *E*. *coli* isolate DNA in *N*. *cinerea* and *A*. *p*. *doriferus* pups at each colony during each breeding season sampled, with percentage of pups that tested positive.

Class 1 integrons were detected at all *N*. *cinerea* and *A*. *p*. *doriferus* colonies sampled (Fig 2). There was a significant difference in the prevalence of class 1 integrons across *A*. *p*. *doriferus* colonies sampled (*χ*^2^_3, 40_ = 58.8, p<0.001). There was no statistically significant difference in prevalence across *N*. *cinerea* colonies (*χ*^2^_2, 11_ = 2.90, p=0.234). The highest number of class 1 integrons (*n*=27) was observed in *A*. *p*. *doriferus* pups at Seal Rocks in 2018. The analysis of prevalence within colonies across breeding seasons revealed a significant difference at Seal Rocks (*χ*^2^_2, 31_ = 40.51, p<0.001) and Seal Bay (*χ*^2^_2, 5_ = 10, p<0.01), but there was no significant difference at Olive Island (*χ*^2^_1, 5_ = 1.8, p=0.179; Fig 2).

### Gene cassette array diversity

DNA sequencing identified five different gene cassette arrays from the 51 positive samples. The majority of the samples (*n*=40) contained gene cassette arrays void of ARGs. Of the samples containing integrons with ARGs, seven contained a single gene cassette and the remaining four arrays each had two gene cassettes (Fig 3).

**Fig 3.**
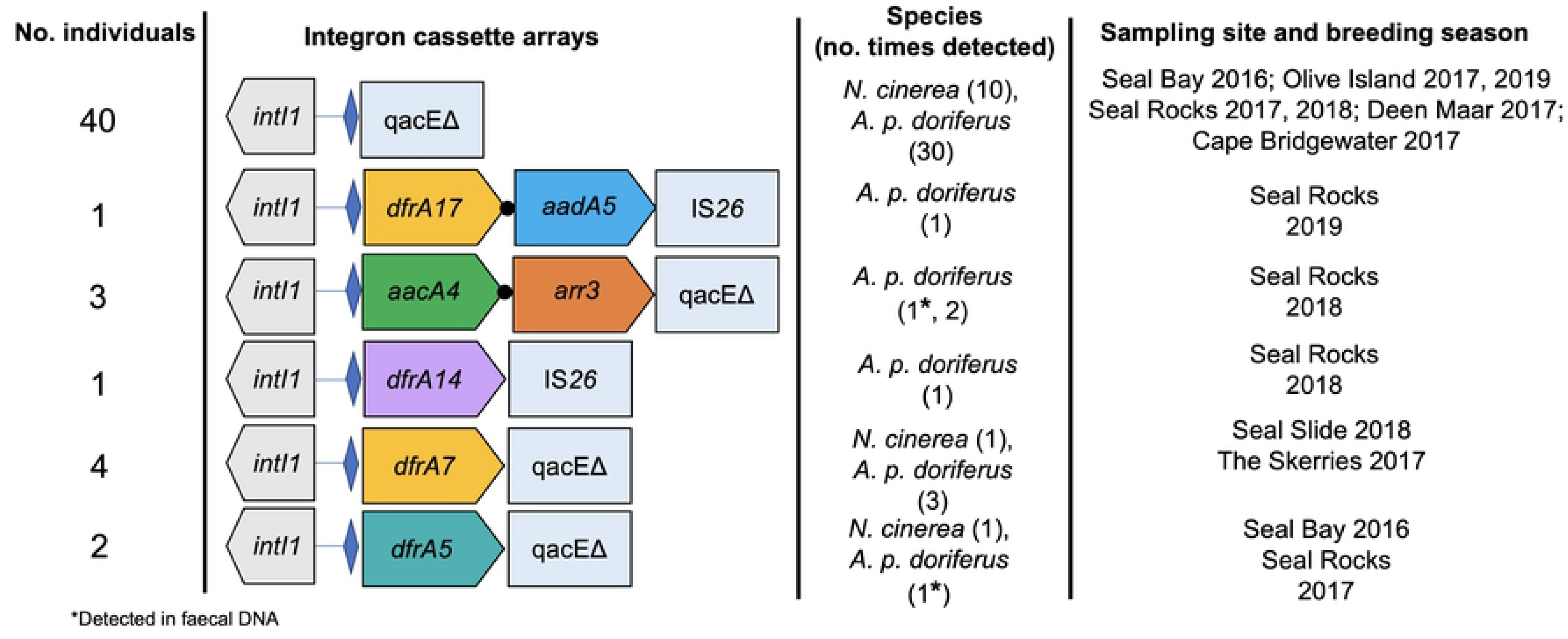
Schematic map of class 1 integron gene cassette arrays identified in *N*. *cinerea* and *A*. *p*. *doriferus* across all sampling sites and breeding seasons. Number of individuals with each array are listed on the left-hand side. Gene cassettes are represented as broad arrows. Blue diamonds represent the primary integron recombination site, *attI1*, where gene cassettes are inserted following acquisition; black circles represent gene cassette recombination site, *attC*. Gene symbols are as follows: *dfrA* genes encode dihydrofolate reductases that confer resistance to trimethoprim; *aacA* genes encode aminoglycoside (6’) acetyltransferases (*aacA*) that confer resistance to aminoglycoside antibiotics; *aadA* genes encode aminoglycoside (3”) adenylyltransferases that confer resistance to streptomycin and spectinomycin; *arr3* genes encode ADP-ribosyl transferases that confer resistance to rifampin; *qac* genes encode efflux pumps that confer resistance to quaternary ammonium compounds; *qacEΔ* and IS*26* represent the 3’ terminus of some the gene cassette arrays depicted.

Class 1 integrons identified in samples from *A*. *p*. *doriferus* were the most diverse, encoding seven different ARGs, while only two types of ARGs were detected in *N*. *cinerea* (Fig 3). The most common cassette array was *dfrA7* (*n*=4), identified in *E*. *coli* isolate DNA from both *N*. *cinerea* and *A*. *p*. *doriferus* pups.

The vast majority (49 of 51) of gene cassette arrays detected in this study contained the typical 3’ conserved segment (*qacEΔ*), whereas, in the remaining two gene cassette arrays, *qacEΔ* was replaced with an IS*26* transposase (Fig 3).

### Essential element and heavy metals and class 1 integron co-selection

There was no significant relationship between the concentrations of Zn (p=0.905; 95% CI 3.67-3.87), Se (p=0.507; 3.00-3.56), As (p=0.446; 0.05-0.07), Hg (p=0.335; 0.05-0.12) or Pb (p=0.937; 0.04-0.10) in whole blood and integron carriage in *A*. *p*. *doriferus* (*n*=52) sampled at Seal Rocks in 2018.

## Discussion

This study identified class 1 integrons in both *E*. *coli* and faecal DNA from free-ranging *N*. *cinerea* and *A*. *p*. *doriferus* pups at seven breeding colonies in Southern Australia, representing the first time ARGs have been detected in *A*. *p*. *doriferus*, and in the faecal microbiota from *N*. *cinerea* pups. The occurrence of class 1 integrons in *A*. *p*. *doriferus* pups is of particular interest given the higher carriage of *intI1* in comparison with the other pinniped species studied.

There was similar class 1 integron abundance and gene cassette diversity across all four *A*. *p*. *doriferus* colonies sampled in 2017. These colonies differ in terms of size, topography, pup production and population density (44) and cover a wide geographical area. The similar *intI1* abundances indicates that these colonies are exposed to similar sources and levels of anthropogenic pollution, however, the number of *intI1* genes detected at Seal Rocks showed considerable change over sampling years (2017-2019), with a significant increase observed in 2018. This increase was not sustained over multiple breeding seasons and further investigation is needed to determine if this increase is due to a gradient of anthropogenic pollution or whether it is an aberrant finding at this colony. Seal Rocks was the only *A*. *p*. *doriferus* colony sampled over multiple breeding seasons limiting our ability to fully assess and compare trends in *intI1* abundance across other *A*. *p*. *doriferus* colonies over time.

The highest number of class 1 integrons was detected in *A*. *p*. *doriferus* pups at Seal Rocks. Class 1 integrons are generally more prevalent in locations closer to urbanised environments that are exposed to higher levels of anthropogenic pollution (13,45). Seal Rocks is located 1.8km from Phillip Island and approximately 7km from the Mornington Peninsula, both of which are densely populated over summer months when pups are born and sampled. Seal Rocks is also located within 150km of Melbourne, Australia’s second largest city (∼5 million people), which could result in continuous exposure of pinnipeds to a higher number of anthropogenic sources of pollution compared to *A*. *p*. *doriferus* pups at the other colonies sampled (for example, Deen Maar Island and Cape Bridgewater are ∼250km and 300km respectively from Melbourne), thereby facilitating the greater acquisition of class 1 integrons.

The abundance of *intI1* is presumed to change rapidly based on environmental factors (13). Rapid changes in environments caused by extreme weather events can also introduce higher levels of runoff and industrial waste into the marine environment and distribute pollutants across wider geographical ranges (46). Environmental conditions at pinniped breeding colonies were not considered as part of this study, however, given the differences in *intI1* abundance observed, the influence of stochastic events on the presence of class 1 integrons would need to be considered when attempting to understand the trends observed in *A*. *p*. *doriferus* pups.

In addition to the higher abundance of *intI1* genes detection in *A*. *p*. *doriferus* pups at Seal Rocks in 2018, was the presence of two IS*26* class 1 integron variants in *E*. *coli* isolates. There are multiple reports of IS*26* class 1 integron variants in *E*. *coli* from animals in Australia, including cows and pigs (14,47,48) and Grey-headed flying foxes (*Pteropus poliocephalus*) (49), as well as in human clinical cases (50). The presence of class 1 integrons with IS*26* variants in *E*. *coli* isolates from free-ranging *A*. *p*. *doriferus* pups provides further evidence to suggest that this colony is exposed to a marine environment that is contaminated by various sources of anthropogenic pollution.

The finding of the *arr3* gene cassette in multiple *A*. *p*. *doriferus* pups at Seal Rocks in 2018 was unexpected. This gene cassette, *arr3*, encodes ADP-ribosyl transferases which potentially confers resistance to rifampicin and has been identified in numerous Gram-negative pathogens (such as *Proteus spp*.) (51,52) and integrons in bacterial isolates from hospital patients (53). The *aacA4*-*arr3*- *qacE*Δ cassette array detected in *A*. *p*. *doriferus* is not widely reported in the literature (53), however, the closest matches on GenBank were predominately detected in *Proteus* spp. and *Klebsiella* spp. (e.g. CP053614, LC549808) isolates from hospital patients and zoo and production animals in China. In this study, the array was detected in faecal DNA and the bacterial carrier remains unknown. As the *arr3* cassette is associated with human and domestic animal pathogens, the presence of this gene cassette in free-ranging pinnipeds provides further evidence to suggest that the Seal Rocks colony is exposed to anthropogenic microbial pollution.

In contrast to *A*. *p*. *doriferus* and *N*. *cinerea*, class 1 integrons were not detected in *A*. *forsteri* pups (*n*=150) sampled over multiple breeding seasons. Over 5000 pups are born at the Cape Gantheaume colony each breeding season (54) and there is a much higher population density compared to Seal Slide, a small *N*. *cinerea* colony <10km away. Despite the similarity in location and higher population density between these two pinniped species a class 1 integron was detected in one of four *N*. *cinerea* sampled pups during the 2018 breeding season. The difference in *intI1* abundance between species is likely multifactorial and further investigation is necessary to better understand the factors that contribute to the acquisition of class 1 integrons in free-ranging pinniped species.

Heavy metals are environmental pollutants that can act as co-selecting agents for antibiotic resistance (16). Heavy metals do not degrade and therefore persist in the environment for long periods of time, maintaining selective pressure for extended periods (18). Furthermore, the abundance of class 1 integrons has been found to correlate with heavy metal concentrations, with previous studies focusing on the relationship between heavy metal pollutants and *intI1* abundance in environmental samples including water and soil (17,55). It has been well established that concentrations of heavy metals are present in free-ranging marine mammals (27,28) and can bioaccumulate in upper trophic predators (26,56). Despite the association between heavy metal concentrations and class 1 integron abundance in environmental samples, this relationship has not previously been investigated in free-ranging wildlife. A significant relationship between the concentrations of ARB carriage and two essential elements and three heavy metals was not seen, although the small sample size for this specific comparison could have limited this analysis. In addition, the current comparison was limited to concentrations of essential and heavy metals in whole blood which reflect recent exposure. The relationship between ARB carriage and heavy metal concentrations in other tissues (for example, liver) could be explored, given the bioaccumulation potential of this tissue when compared to whole blood (57).

Free-ranging wildlife are exposed to differing environmental conditions, selective pressures and exposure to ARB that likely drive the acquisition of ARGs (58). Pinniped pups sampled as part of this study were less than two months of age and confined to the breeding colonies. As such, the only potential source of exposure to bacteria and ARGs for pups are those present in the breeding colony environment, which could be contaminated by wastewater run-off, faecal contamination from other wildlife species (including sea birds) and juvenile and adult pinnipeds that inhabit these colonies (32,59). Evidence suggests that the prevalence of ARB in free-ranging wildlife is influenced by exposure to anthropogenic pollution and environmental contamination (58), with the latter varying depending on habitat occupation and behaviours exhibited by wildlife species (58). For example, foraging is a behaviour that can increase the likelihood of wildlife species being exposed to ARB (3). The pinniped species studied herein have differing foraging strategies. While *N*. *cinerea* and *A*. *p*. *doriferus* are benthic foragers (60–63), *A*. *forsteri* are pelagic foragers (64,65). In some aquatic species, differences in toxicant accumulation between benthic and pelagic feeders has been observed (57), thus differences in foraging behaviours could lead to exposure of higher levels of pollutants and increased ARB acquisition in benthic feeding pinniped species. Interactions with other species in ecosystems is another aspect to consider when attempting to understand the transfer of AMR. The presence of other species at pinniped breeding colonies such as sea birds, known carriers of numerous ARGs (59), could also influence the carriage of AMR in free-ranging pinnipeds. Sampling both environmental substrates and other wildlife species present within study areas is required to gain a greater understanding of the source of AMR and the transfer of ARGs in free-ranging wildlife species.

## Conclusion

This study detected bacteria carrying diverse gene cassettes encoding resistance to multiple classes of antibiotics in two species of free-ranging pinniped pups in Australia. The detection of class 1 integrons, mobile genetic elements that have been identified as useful indicators of antimicrobial pollution, suggests these populations are exposed to anthropogenic pollution. Furthermore, the detection of *E*. *coli* carrying IS*26* class 1 integron variants in free-ranging pinniped pups indicates that these isolates originated from domestic animals and/or humans. Further investigation to better understand how antibiotic resistant bacteria are being acquired by free-ranging pinniped pups is critical and could be used as an additional mechanism to monitor anthropogenic pollution in marine ecosystems. Ongoing monitoring of antibiotic resistant bacteria in these species will also assist in understanding the role of increasing anthropogenic pollution on the long-term survival of these marine sentinel species.

## Acknowledgements

We thank staff at Phillip Island Nature Parks (PINP), Victoria and staff at Seal Bay, Kangaroo Island, Department for Environment and Water (DEW), South Australia particularly Melanie Stonnill, for field assistance and logistical support; Simon Goldsworthy and South Australian Research and Development Institute (SARDI) for field assistance. Sample collection was made possible through the collaborative support of DEW, PINP and SARDI. We would like to thank Scott Lindsay, Shannon Taylor and Matthew Gray for assistance in sample collection; Victorian Fisheries Authorities, T-cat Charters, Seatec Marine services and Darren Guidera for marine charters. We thank Robert McQuilty, Royal Prince Alfred Hospital NSW, for his assistance in trace and heavy metal analysis.

